# Developmental, neuroanatomical and cellular expression of genes causing dystonia

**DOI:** 10.1101/2025.04.13.648593

**Authors:** Darren Cameron, Nicholas E Clifton, Daniel Cabezas de la Fuente, Peter Holmans, Nicholas J. Bray, Kathryn J Peall

## Abstract

**Objective:** Dystonia is one of the most common forms of movement disorder with >50 genes identified as causative. However, an understanding of which developmental stages, brain regions and cell types are most relevant is crucial for developing relevant disease models and therapeutics. One approach is to examine the timing and anatomical expression of dystonia-causing genes, on the assumption that deleterious variants have a greater impact where higher levels of expression are observed.

**Methods:** We investigated the expression patterns of 44 dystonia-causing genes across two bulk- and two single-nuclei RNA-sequencing datasets, derived from prenatal and postnatal human brain tissue.

**Results:** Dystonia genes were most strongly enriched in those with higher expression in the striatum, cerebral cortex, hippocampus, amygdala and substantia nigra, and for higher postnatal expression. Individual genes exhibiting differences in expression across adult brain regions include *SQSTM1, SGCE, KMT2B, PRKRA, YY1, DNAJC12, KCNA1, CACNA1A* (highest expression in cerebellum), *ADCY5, GNAL, ANO3* (highest expression in striatum), *RHOBTB2, FOXG1* (highest expression in cerebral cortex). Single-nuclei RNA-sequencing data analyses from human frontal cortex, striatum and cerebellum indicated that dystonia genes are predominantly expressed in neurons (both glutamatergic and GABAergic), rather than glia. Gene Ontology analysis showed prominent enrichment in biological processes such as *dopamine biosynthetic and metabolic processes*, and in the cellular components *axons, presynapse* and *neuron projection*.

**Interpretation:** These analyses provide important insights into the anatomical, developmental and cellular expression patterns of dystonia-causing genes, potentially guiding the development of disease-relevant models and improving the timing and targeting of future therapeutic interventions.

## Introduction

Dystonia is a hyperkinetic movement disorder estimated to affect 120/100,000 people, and is characterized by involuntary, repetitive, or sustained muscle contractions resulting in abnormal posturing, pain, functional impairment, and reduced quality of life.^1,2^ It can arise as a primary disorder, or part of a broader neurodevelopmental or neurodegenerative condition, with both genetic and idiopathic forms, and a spectrum of motor presentations.^3,4^ Over the past two decades, and accelerated with the advancement of next-generation sequencing technologies, more than 50 genes have now been linked to the dystonia clinical phenotype.^5^

The identification of monogenic causes of dystonia has facilitated the study of genetically homogenous clinical cohorts and the development of model systems across mammalian, invertebrate and cellular models, enabling investigation of the underlying pathophysiological mechanisms in dystonia.^6,7^ Collectively, studies suggest that dystonia is driven by disruption to neural networks, with particular involvement of the cortico-basal ganglia-thalamo-cerebellar network. *In vivo* human imaging studies have shown disrupted network-based cerebral motor control,^8^ with reduced functional connectivity between the basal ganglia, sensorimotor and frontoparietal regions.^9^ Structural imaging studies in individuals harbouring *TOR1A* and *THAP1* mutations have revealed white matter changes in the sensorimotor cortex and outward cerebellar projections,^10,11^ with similar disruption observed in the sensorimotor cortex, caudate and putamen in cohorts with *ATP1A3* and *SGCE* mutations.^12,13^ Animal models have implicated disrupted striatal GABA transmission^14^ and cerebellar Purkinje cell abnormalities,^15^ while stem cell models have demonstrated functional hyperexcitability and defects in synaptic plasticity and cellular stress response pathways.^16^ Furthermore, an unbiased genetic systems biology approach highlighted key roles for both the cerebral cortex and basal ganglia in dystonia pathogenesis.^17^

Pathway analyses have provided additional insights. Recent RNA-sequencing analyses in brain tissue derived from *Tor1a* knock-in embryonic mice (E18), identified”eIF2(alpha) Signalling” and “Cell Cycle checkpoint regulation” as the highest ranked dysregulated pathways in homozygous lines, and “Axonal Guidance Signalling” and “Actin Cytoskeleton Signalling” in their heterozygous counterparts.^18^ Moreover, genetic engineering of eight distinct *THAP1* variants into a common induced pluripotent stem cell line, followed by bulk RNA-sequencing, revealed common pathways related to neurodevelopment, lysosomal lipid metabolism and myelin.^19^

The few dystonia studies involving human brain tissue undertaken to date have focussed exclusively on adult-derived samples and typically examined the expression of single dystonia-causing genes.^20,21^ No previous work has comprehensively explored the expression patterns of multiple dystonia-causing genes across the human lifespan, in different brain regions, or in both neuronal and non-neuronal cell types. However, the increasing availability of large bulk- and single-nuclei RNA-sequencing datasets now makes it possible to conduct both cross-sectional and longitudinal analyses. In this study, we leverage four independent human brain gene expression datasets, spanning eight developmental stages, utilising both bulk- and single nuclei RNA-sequencing data.

Gaining a deeper understanding of the temporal, regional and cellular expression of dystonia-causing genes will not only help identify therapeutic targets but also aid in pinpointing the optimal developmental window for therapeutic intervention to achieve clinical benefit.

## Materials and Methods

### Monogenetic causes of a dystonia-predominant phenotype

This study focuses on monogenic inherited movement disorders where dystonia is the predominant motor phenotype. A list of 44 dystonia-causing genes (Supplementary Table 1) was curated from previously published gene lists linked to monogenic forms of dystonia, including isolated, combined and complex forms.^22,23^ Genes were excluded if linked to neurodegeneration with brain iron accumulation (NBIA), for example *ATP7B, PANK2, CP, PLA2G6, SLC30A10, SLC39A14* and metabolic disorders (*GCDH, HPRT, GLB1, PTS, QDPR*). In addition, those reported in only a limited number of cases to date (*DACF17, IRF2BPL, ADAR1*) or coupled with yet unconfirmed DYT genes (*CIZ1, COL6A3, RELN*) were also not included.

### RNA-Seq datasets

Two bulk and two single nuclei RNA-Seq datasets were analysed in this study, initially using the bulk RNAseq datasets before moving onto the single cell nuclei analysis to gain more specific cellular understanding.

### Bulk RNA-Seq datasets and analysis

#### BrainSpan dataset

Human brain RNA-Seq data were accessed from the BrainSpan repository (https://www.brainspan.org/static/download.html).^24^ The gene expression matrix contained RPKM values derived from 524 independent brain tissue dissections across 42 donors, aged 8 weeks post-conception to 40 years. These dissections were assigned to 31 unique age annotations which we aggregated into 7 developmental stages (early fetal, mid fetal, late fetal, infancy, childhood, adolescence, adulthood, Supplementary Table 2). Samples with ambiguous metadata and low RNA integrity (RIN < 7) were excluded, leaving 39 donors and 447 tissue dissections for downstream analyses.

To assess gene expression differences in dystonia-causing genes across developmental stages, a two-step linear modelling approach was applied. In step one, gene expression values were Yeo-Johnson transformed, and separate linear models were fitted for each dystonia gene to adjust for covariates RIN, ethnicity, and sex:

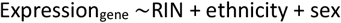

The residuals of each model, representing each dystonia gene’s expression after adjusting for covariate effects, were extracted and the residuals for all genes were aggregated for the next step. In step two, to assess dystonia gene expression across developmental stages, eight linear models were fitted to the aggregated residuals using the following model:

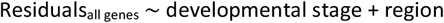

Here, “developmental stage” was a binary variable indicating whether the residuals corresponded to a specific developmental stage (e.g., Early Fetal) or not, and “region” was a categorical variable encompassing all brain regions. This analysis was performed independently for each developmental stage to assess its contribution to variation in dystonia gene expression relative to all other developmental stages, whilst accounting for regional effects.

Effect sizes and p-values were extracted from the step two models, and Benjamini-Hochberg correction (FDR)-adjusted p-values were calculated. For visualization, −log_10_(FDR-adjusted p-values) were plotted. Significant effects (P_FDR_ < 0.05) are highlighted in blue.

#### FUMA analyses

The GENE2FUNC component of the FUMA2 online tool (http://fuma.ctglab.nl) was used to assess tissue-specific expression patterns of dystonia genes based on GTEx v8 RNA-seq data,^25^ which includes RPKM values for 54 tissue types and 22,146 genes.

For the tissue-specific expression analyses, dystonia-causing genes were evaluated at two levels in GTEx v8 (54 detailed tissue types and 30 general tissue categories), using FUMA’s pre-defined differentially expressed (DEG) gene sets, which represent genes with significantly higher expression in a single tissue type compared to all others. Enrichment of dystonia-causing genes within each tissue-specific DEG set was tested using a hypergeometric test, with a background set restricted to protein coding genes. Results were visualised in bar charts, with a significance level of Bonferroni corrected P < 0.05. Hierarchical clustering was performed on the average log2-transformed RPKM expression values of the 44 dystonia-causing genes to identify dystonia genes with similar expression patterns across the 54 GTEx v8 detailed tissue types.

To explore the biological functions, pathways and functional categories associated within the dystonia-causing gene set, hypergeometric tests were performed to assess enrichment of dystonia genes in curated gene sets from MsigDB and WikiPathways. Results were visualised in bar charts showing only significant pathways (P_FDR_ < 0.05), including enrichment P-values, proportions of overlapping genes, and the extent of overlap with each gene set. An FDR correction was applied for multiple testing, with a significance threshold of P_FDR_ < 0.05.

### Single nuclei RNA-Seq data preparation and analysis

#### Prenatal brain data

We used single nuclei RNA-Seq (snRNA-Seq) data that we have previously generated ^26^ from dissected regions of the human fetal brain (3 samples aged 14-15 post-conception weeks) to assess dystonia gene expression in individual prenatal brain cell populations. A detailed description of the quality control and data processing measures is provided in that paper ^26^. For this study, we focused on data generated from 3 fetal brain regions (frontal cortex, cerebellum and the ganglionic eminences [the latter generating inhibitory neurons of the cerebral cortex and striatum]) corresponding to those regions of the adult brain that we also analysed using snRNA-Seq data. Violin plots were used to display the normalized expression scores of the 44 dystonia-associated genes across different cell types in each region.

#### Postnatal brain data

snRNA-Seq data from adult human brain were downloaded from the CELLxGENE repository (https://cellxgene.cziscience.com/collections/283d65eb-dd53-496d-adb7-7570c7caa443).^27^ In the original study by Siletti and colleagues, 100 tissue dissections from 3 human donors aged between 18-68 years were performed across 10 brain regions. For this study, data from 3 brain regions were accessed, encompassing 13 independent dissections (frontal cortex=8, striatum=2, cerebellum=3 (Supplementary Table 3). A separate R object was downloaded for each dissection that contained a matrix of raw unique molecular identifier (UMI) counts and associated metadata. All downstream processing was carried out using the scRNAseq software suite Seurat 5.0.2 using default settings unless otherwise stated.

For each region, dissection-specific R objects were combined into a single Seurat object, and the data split by sequencing lane, referred to henceforth as ‘sample’, such that quality control and data integration were carried out on a per sample basis. In addition to the quality control measures applied to the data in the original study,^27^ outliers were identified, and cells excluded if their log-transformed library sizes were >3 median absolute deviations above the median cell UMI count per sample, or if >5% of their UMIs mapped to mitochondrial or ribosomal genes. *MALAT1*, genes mapping to the mitochondrial genome and genes expressed in <3 cells were also removed. Final dimensions of the gene expression matrices (genes × cells) for each region were: 26,795 × 331,113 (frontal cortex); 30,138 × 144,380 (cerebellum) and 27,379 × 66,782 (striatum).

Due to the size of the adult datasets, a sketch object of approximately 60K cells was generated for each region. A sketch object is designed to make the analysis of large datasets less memory intensive by extracting a representative subset of cells for each region that preserves their underlying data structure. First, the data were normalised and the top 2000 most variable genes identified, *Seurat::SketchData* was then run to generate a sketch object for each region. Variable genes (again the top 2000) were identified for the sketched data, the data were scaled, and a principal components analysis was run, retaining the first 30 principal components for the striatum and cerebellum, and 50 for the frontal cortex. The data integration software Harmony was used to correct sample specific batch effects, cells were clustered with a resolution setting of 0.1, and the clusters were visualised in two-dimensional space via Uniform Manifold Approximation and Projection (UMAP). Cell type identity was assigned to each cluster based on expression of known canonical markers (Supplementary Figures 1-3). Finally, dimensional reductions and cluster labels identified on the sketched data for each region were projected onto the entire dataset, and normalised expression scores for the 44 dystonia genes across cell types were visualised using violin plots.

**Figure 1.**
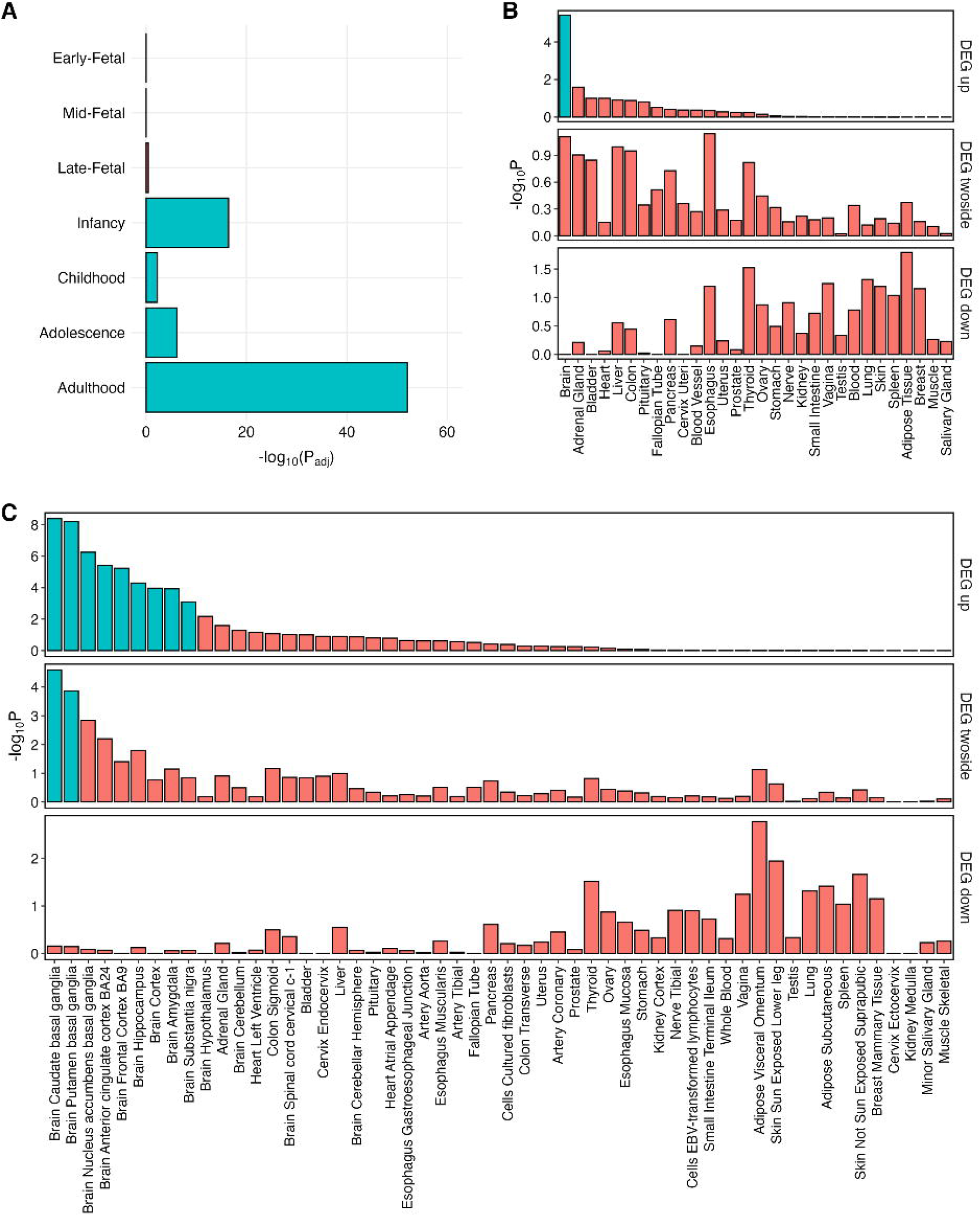
Temporal and tissue expression of dystonia causing genes. A. **BrainSpan dataset**. log_10_(FDR-adjusted p-values) across seven developmental stages. Positive or negative signs correspond to the direction of the effect estimates. Significant effects (P_FDR_ < 0.05) are highlighted in green. **B: GTEx Dataset**. Differential Gene Expression of selected dystonia genes in the GTEx v8 30 general tissue types. C: **GTEx Dataset**. Differential Gene Expression of all dystonia genes in the GTEx v8 54 general tissue types. Significantly enriched (p<0.05, following Benjamini-Hochberg (FDR) correction for multiple testing) differentially expressed genes (DEG) are highlighted in red.

#### Cellular enrichment of dystonia genes

To investigate the cellular specificity of dystonia gene expression, we applied the Expression Weighted Cell Type Enrichment (EWCE) method to the pre- and post-natal snRNA-seq datasets. These analyses aimed to determine whether the 44 dystonia genes collectively exhibit enriched expression in specific cell types relative to all other cell types within each brain region. For each brain region, raw UMI counts were normalized using SCTransform, and gene expression specificity scores (GESS) were calculated for each gene in each cell type using EWCE. GESS values (range 0-1), indicate the proportion of a gene’s expression attributed to a single cell population relative to its total expression across all cell populations within that region. To assess enrichment, we conducted a bootstrap-based test with 10,000 iterations, comparing the specificity scores of the 44 dystonia genes against those of random gene sets of equal size, drawn from all expressed genes in the brain region. Statistical significance was determined using an FDR correction, with enrichments considered significant at FDR<0.05.

### Predicted protein-protein interactions

The STRING database (https://string-db.org/) was used to identify predicted protein-protein interactions in implicated neurons of the adult striatum, frontal cortex and cerebellum based on data derived from experimental and database evidence.

## Results

### Increased dystonia gene expression in the postnatal period

Using the bulk RNA-Seq BrainSpan dataset, we assessed differences in dystonia gene expression in human brain across 7 developmental stages. We focussed on identifying patterns of expression by modelling expression at each developmental stage independently, adjusting for brain region effects. The analyses revealed significantly higher relative dystonia gene expression in all four postnatal stages (infancy, childhood, adolescence and adulthood) when compared individually to the remaining developmental stages (Figure 1A, Supplementary Table 4). The most pronounced increases were observed in adulthood (P = 8.43 × 10^−53^) and infancy (P = 3.65 × 10^−17^), with more moderate relative increases in adolescence (P = 7.89 × 10^−7^) and childhood (P = 5.73 × 10^−3^).

### Enrichment of dystonia gene expression in specific adult brain regions

Bulk RNA-Seq GTEx data was used for dystonia gene expression analysis in adult human tissue. Here, dystonia-causing genes were significantly enriched in those upregulated in brain tissue, compared to non-brain tissue (P = 3.73 × 10^−6^, Figure 1B). Analysis of individual brain regions, revealed an upregulation of dystonia genes in constituent subregions of the striatum (caudate (P = 4.07 × 10^−9^), putamen (P = 5.99 × 10^−9^), nucleus accumbens (P = 5.60 × 10^−7^)), as well as anterior cingulate cortex (P= 3.85 × 10^−6^), frontal cortex (P = 5.79 × 10^−6^), hippocampus (P = 5.04 × 10^−5^), cerebral cortex (P = 1.07 × 10^−4^), amygdala (P = 1.18 × 10^−4^) and substantia nigra (P = 8.12 × 10^−4^) (Figure 1C).

### Expression of individual dystonia genes in adult brain regions

We next explored the neuroanatomical expression of individual dystonia-causing genes using GTEx data. Prominent expression was observed in the adult cerebellum, striatum, cortex, amygdala, and hippocampus, with several genes demonstrating high levels of expression across all these regions (*SLC2A1, ACTB, BCAP31, GNB1, PNKD, GNAO1, ATP1A3, TUBB4A, HPCA, TMEM151A*, Figure 2). Other dystonia genes displayed more region-specific patterns, including *SQSTM1, SGCE, KMT2B, PRKRA, VPS16, YY1, DNAJC12, KCNA1, CACNA1A* (highest expression in cerebellum), *ADCY5, GNAL, ANO3* (highest expression in striatum) and *RHOBTB2* and *FOXG1* (highest expression in cerebral cortex). These results highlight the region-specific regulation of individual dystonia genes, and the potential importance of the cerebellum for mediating effects of certain gene disruptions.

**Figure 2.**
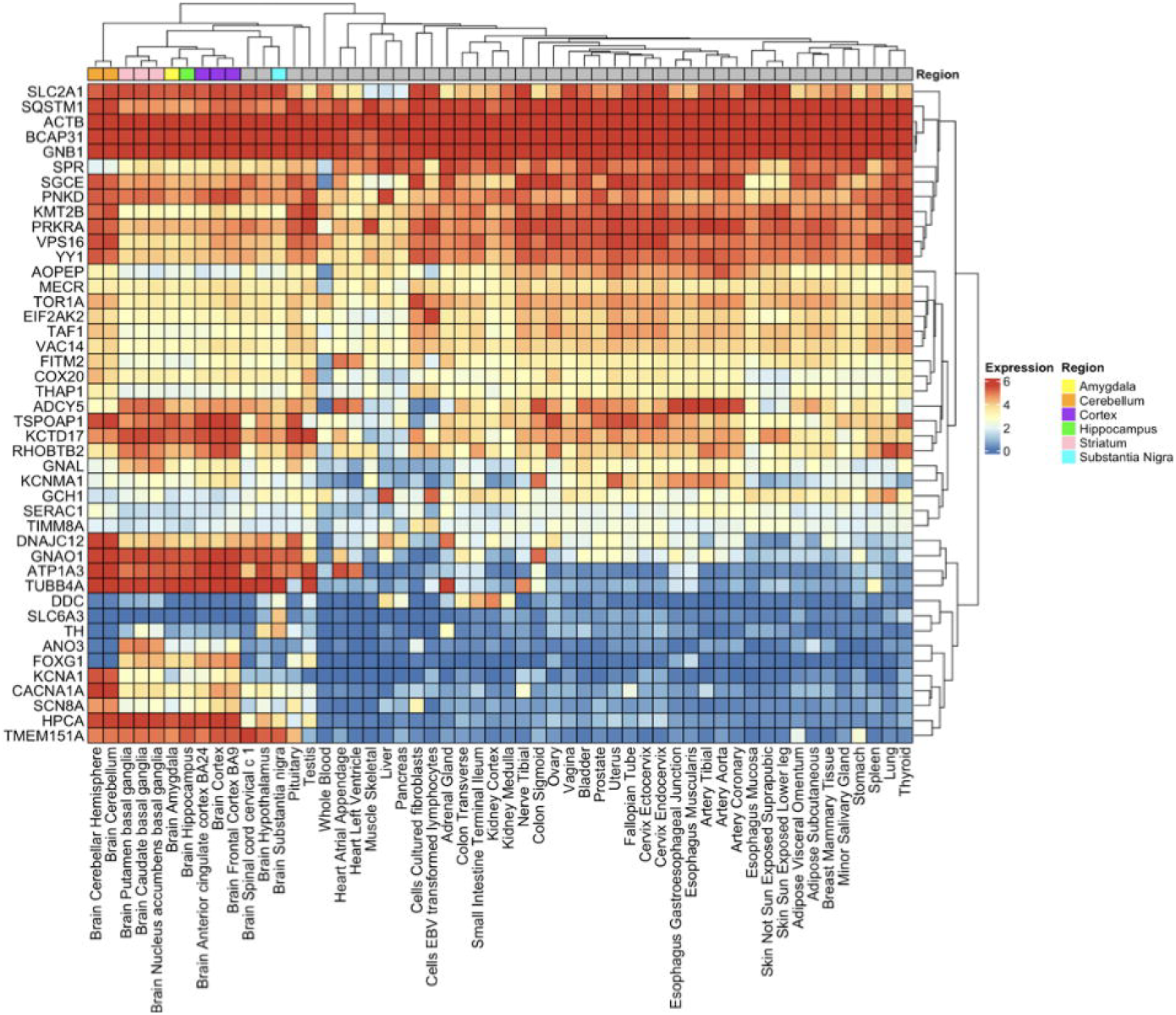
Variation in tissue expression across individual dystonia genes. **GTEx dataset**. Gene expression heatmap of the average log2 transformed RPKM expression values per tissue type, using the GTEx v8 54 general tissue type dataset. Darker red indicates higher expression compared to the darker blue colour indicating lower or no expression. Clustered tissue types include Amygdala (Yellow), Cerebellum (Orange), Cortex (Purple), Hippocampus (Green), Striatum (Pink), Substantia Nigra (Blue)

**Figure 3.**
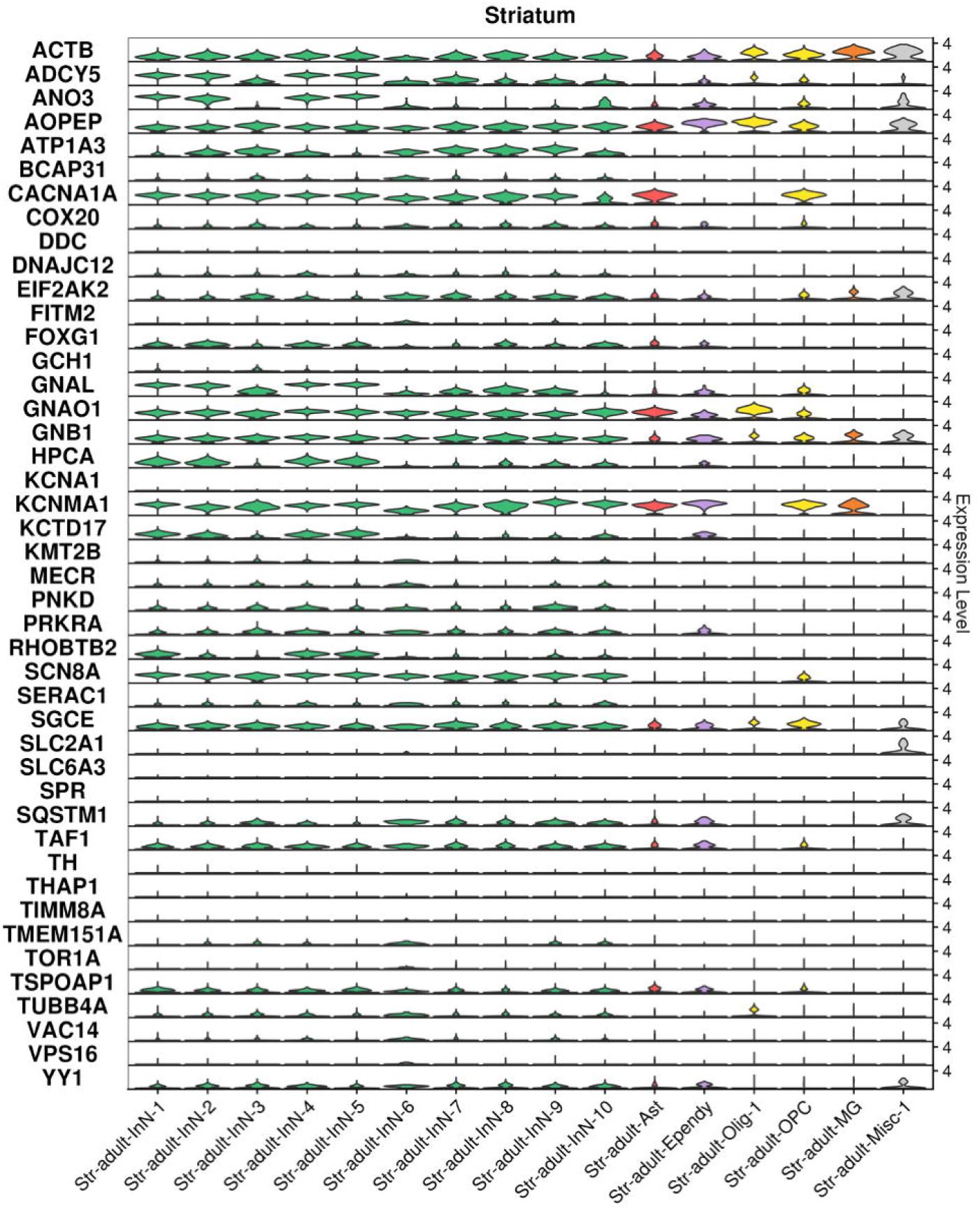
Cellular expression of individual dystonia genes in adult striatal tissue. Violin plots demonstrating expression levels of individual dystonia genes in adult Striatal tissue. Ast: Astrocytes, Ependy: Ependymal cells, InN: inhibitory neuron, MG: microglia, Misc: Miscellaneous, Olig:Oligodendrocyte, OPC: oligodendrocyte precursor cell. Data derived from previously reported dataset.^27^

### Expression of dystonia genes in individual human neural cell populations

Given observed differences in individual dystonia gene expression across the frontal cortex, striatum and cerebellum, we next explored the cellular specificity of these genes using snRNA-Seq data from corresponding regions of the fetal ^26^ and adult ^27^ human brain.

#### Expression of dystonia genes in cell populations of the fetal brain

Consistent with our findings using the BrainSpan dataset (Figure 1A), fewer dystonia genes were expressed, and at lower levels, in fetal brain cells compared with adult brain cells (Supplementary Figure 4). Using EWCE, we observed no significant (FDR < 0.05) enrichment of dystonia gene expression in any individual cell population within analysed regions of the fetal brain (Supplementary Figure 5). Of the dystonia genes analysed, *ACTB, FOXG1, GNAO1* and *GNB1* were the most strongly expressed in fetal brain cell populations, with all displaying broad expression across neuronal and non-neuronal cells (the latter including radial glia, microglia and endothelial cells).

**Figure 4.**
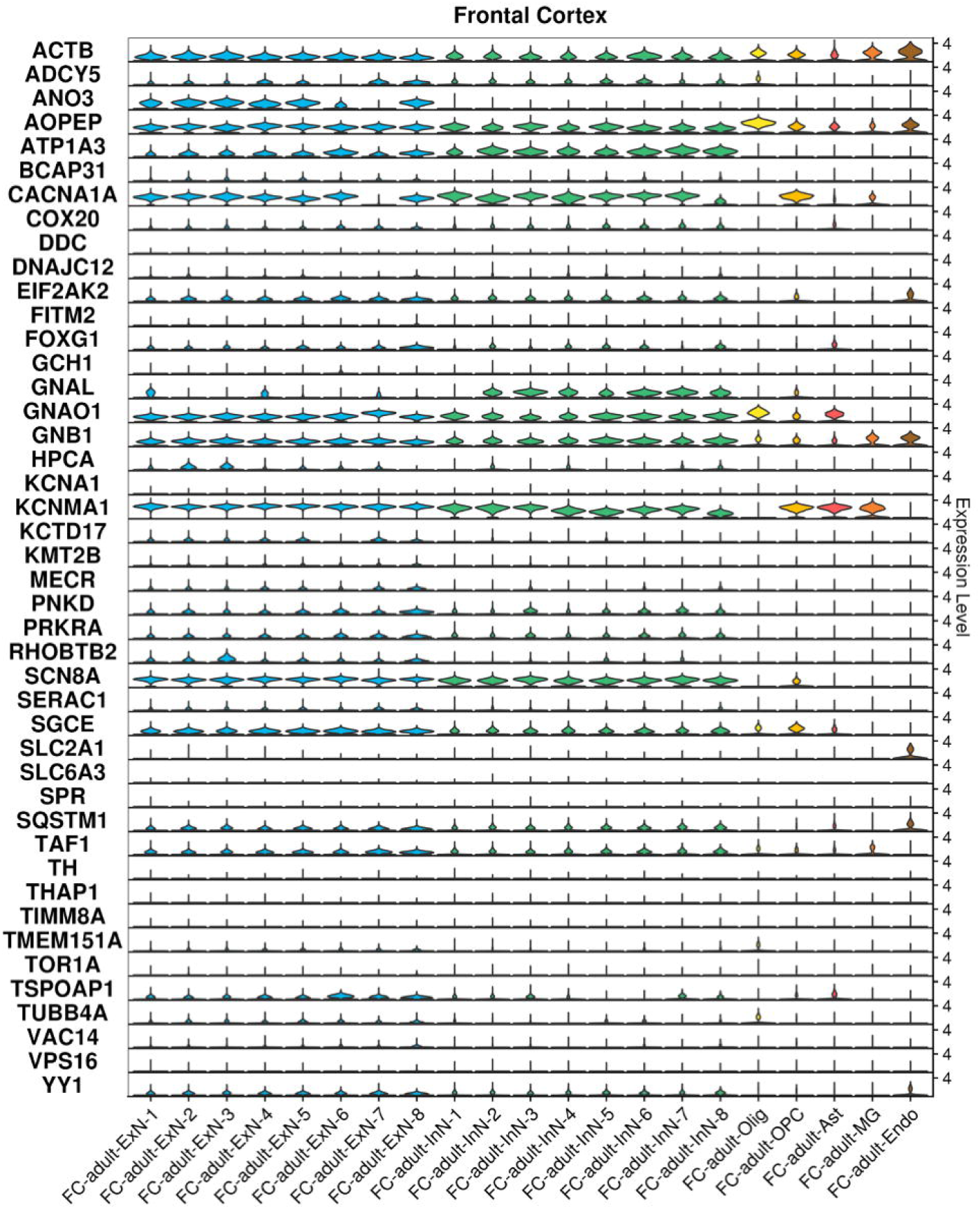
Cellular expression of individual dystonia in genes in adult frontal cortical tissue. Violin plots demonstrating expression levels of individual dystonia genes in adult Frontal Cortex. Ast: Astrocytes, Endo: Endothelial cells, ExN: excitatory neuron, InN: inhibitory neuron, MG: microglia, Olig: Oligodendrocyte, OPC: oligodendrocyte precursor cell. Data derived from previously reported dataset.^27^

**Figure 5.**
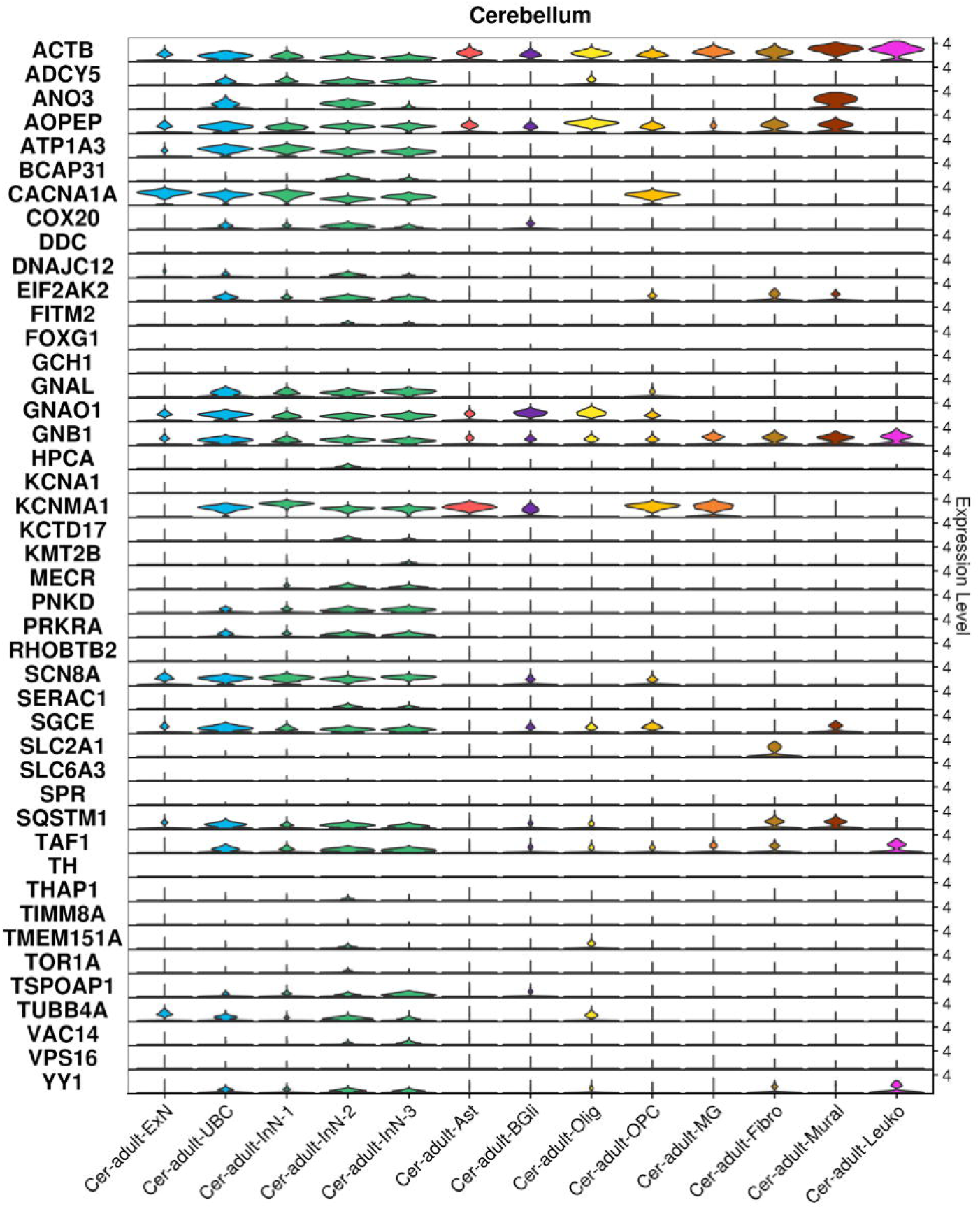
Cellular expression of individual dystonia in genes in adult cerebellar tissue. Violin plots demonstrating expression levels of individual dystonia genes in adult Cerebellar tissue. Ast: Astrocytes, BGli: Bergmann glial cells, Cer: Cerebellum, ExN: excitatory neuron, Fibro: Fibroblast, InN: inhibitory neuron, Leuko: Leukocyte, MG: microglia, Mural: Mural cells, Olig: Oligodendrocyte, OPC: oligodendrocyte precursor cell, UBC: Unipolar Brush cells. Data derived from previously reported dataset.^27^

*ACTB, AOPEP, GNAO1, GNB1, SCN8A, SGCE* and *TAF1* expression was observed in cells across all three fetal brain regions, while *KCNMA1* and *CACNA1A* were more highly expressed across the fetal frontal cortex and cerebellum, and *FOXG1* in the frontal cortex and ganglionic eminences.

#### Expression of dystonia genes in cell populations of the adult striatum

In line with our finding of generally higher expression of dystonia genes in the adult striatum (Figure 1C), the majority of dystonia genes were found to be expressed in striatal neuron populations which consist primarily of GABAergic medium spiny neurons (MSNs) (Figure 3, Supplementary Table 5). For most dystonia genes, expression was detected across both dopamine D1 receptor-expressing (e.g. Str-adult-InN-1, Str-adult-InN-2, Str-adult-InN-3) and dopamine D2 receptor-expressing (e.g. Str-adult-InN-4, Str-adult-InN-5, Str-adult-InN-6) MSN populations. *ACTB, AOPEP, CACNA1A, GNAO1, GNB1, KCNMA1* and *SGCE* were also prominently expressed in non-neuronal cell populations, with *KCNMA1, CACNA1A, GNAO1* and *AOPEP* highly expressed in astrocytes; *AOPEP, ACTB, GNAO1* in oligodendrocytes; *KCNMA1, CACNA1A, ACTB, SGCE* in oligodendrocyte precursor cells (OPCs), and *ACTB* and *KCNMA1* in microglia. Considerably lower expression of *DDC, GCH1, KCNA1, SLC2A1, SLC6A3, SPR, TH, THAP1, TIMM8A, TOR1A, VPS16* was observed across all cell populations of the striatum. However, our EWCE analysis indicated that no single cell population of the striatum was significantly (FDR < 0.05) enriched for dystonia gene expression above others (Supplementary Figure 6A).

**Figure 6.**
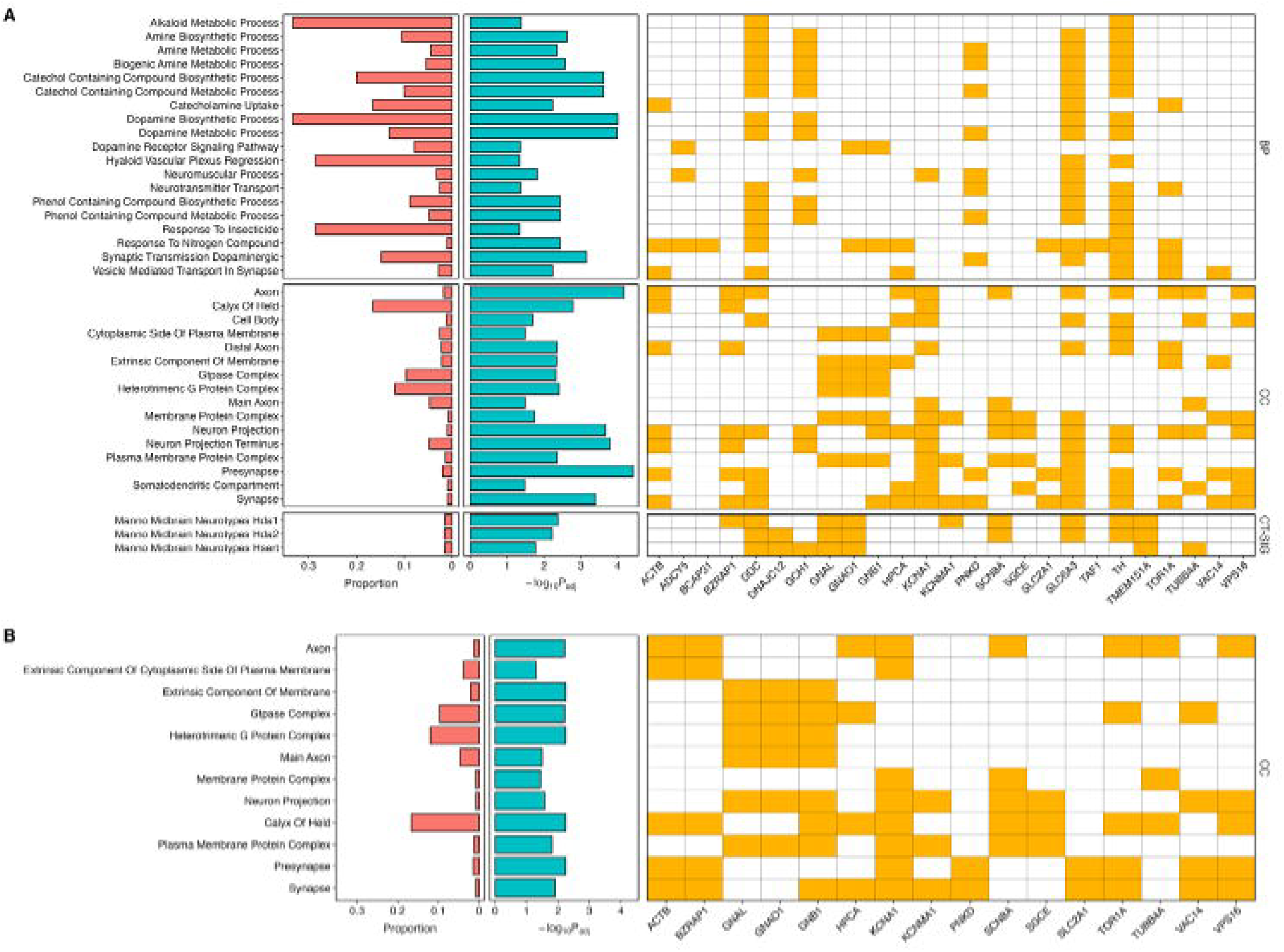
Gene Ontological Analysis in adult brain tissue. Gene set enrichment analysis highlighting only gene sets with an adjusted p-value <0.05 following Benjamini-Hochberg (FDR) correction for multiple testing. Each schematic demonstrates the proportion of overlapping genes in the gene set (red) and enrichment adjusted p-value (blue). These span GO biological processes (MsigDB c5), (B) Cell type signatures (MsigDB c8), (C) GO cellular components (MsigDB c5). Figure A includes all statistically significant (adjusted p<0.05) pathways identified using the whole dystonia gene set (n=44 genes). Figure B demonstrates the statistically significant pathways identified following removal of dystonia genes involved in the dopamine metabolic pathway (*GCH1, TH, SPR, DDC, SLC6A3*). Manno Midbrain Neurotypes Hda1: D1 receptor midbrain dopaminergic neurons, Manno Midbrain Neurotypes Hda2: D2 receptor midbrain dopaminergic neurons, Manno Midbrain Neurotypes Hsert: Midbrain Serotonergic Neurons.

#### Expression of dystonia genes in cell populations of the adult frontal cortex

*KCNMA1, SCN8A, CACNA1A, GNAO1, AOPEP, ATP1A3, ACTB, GNB1, SGCE, EIF2AK2, SQSTM1* and *TAF1* were found to be prominently expressed across excitatory (i.e. glutamatergic) and inhibitory (i.e. GABAergic) cortical neurons (Figure 4, Supplementary Table 6). By contrast, *ANO3, FOXG1, MECR, RHOBTB2, TSPOAP1* and *YY1* were more highly expressed in excitatory cortical neurons, while *GNAL* was more highly expressed in their inhibitory counterparts. *KCNMA1, AOPEP, GNAO1, ACTB, CACNA1A, GNB1* and *SGCE* were additionally expressed in several non-neuronal cell populations, including astrocytes (*KCNMA1, GNAO1, AOPEP)* oligodendrocytes (*AOPEP, ACTB, GNAO1*), OPC (*KCNMA1, CACNA1A, ACTB, AOPEP, SGCE*) and microglia (*KCNMA1, ACTB, GNB1*). No individual cell population of the adult frontal cortex was selectively enriched for dystonia gene expression at the FDR < 0.05 threshold (Supplementary Figure 6B).

#### Expression of dystonia genes in cell populations of the adult cerebellum

The adult cerebellum contains a variety of neuronal populations not found in the other analysed regions of the adult human brain, including the inhibitory (GABAergic) Purkinje cells and excitatory (glutamatergic) unipolar brush cells. Several dystonia genes (*CACNA1A, KCNMA1, AOPEP, GNAO1, SCN8A, ACTB*) were most highly expressed in unipolar brush cells, while *CACNA1A, ATP1A3, AOPEP, SCN8A, GNAL, GNAO1, GNB1* and *SGCE* demonstrated higher expression in Purkinje cells (Figure 5, Supplementary Table 7). Dystonia gene expression in non-neuronal cells included astrocytes (*KCNMA1, ACTB, AOPEP*), OPCs and oligodendrocytes (*AOPEP, KCNMA1, ACTB, GNAO1, GNB1* and *SGCE*), and microglia (*KCNMA1, ACTB, GNB1*). No individual cell population of the adult cerebellum was enriched for dystonia gene expression above others at the FDR < 0.05 threshold (Supplementary Figure 6C).

### Dystonia genes involved in neuronal and synaptic structure and function

Gene set enrichment analyses were undertaken to identify biological processes and cellular substructures in which dystonia genes are over-represented. This revealed significant enrichment across several GO biological processes, cellular components and cell type signatures (P_adj_ < 0.05), primarily related to neurons. Dystonia genes were enriched for annotation to the biological processes Dopamine Biosynthetic (*DDC, GCH1, SLC6A3, TH*) and Metabolic (*DDC, GCH1*, PNKD, *SLC6A3, TH*) Processes, and Synaptic Transmission Dopaminergic (*PNKD, SLC6A3, TOR1A*) (Figure 6A). In cell type signatures, there was enrichment in midbrain monoaminergic neurons and serotonergic neurons, although with a low proportion of overlapping genes. GO cellular pathway analysis further highlighted pathways related to axonal and synaptic structure, including *Presynapse* (*ACTB, BZRAP1, KNA1, PNKD, SLC2A1, SLC6A3, TH, TOR1A, VAC14, VPS16*), *Axon* (*ACTB, BZRAP1, DDC, HPCA, KCNA1, SCN8A, SLC6A3, TH, TOR1A, TUBB4A, VPS16*) and *Neuron Projection Terminus* (*ACTB, BZRAP1, GCH1, KCNA1, SLC6A3, TH*). Given the prominence of dopaminergic pathways and the potential influence of dystonia genes that play a direct role in dopaminergic metabolism and neurotransmission, we reanalysed the data excluding these genes (*GCH1, TH, SPR, DDC, SLC6A3*) (Figure 6B). Even after these exclusions, significant enrichment persisted in pathways such as *Main Axon (GNAL, GNAO1), Axon (HPCA, KCNA1, PRRT2, SCN8A, TOR1A, TUBB4A, VPS16)* and *Heterotrimeric G-Protein Complex (GNAL, GNAO1, HPCA, TOR1A)*, underscoring the central role for neuronal structure and signalling in dystonia.

### Dystonia protein interaction networks are observed across affected brain regions

Seeking to determine potential dystonia protein networks, all dystonia genes with median expression values >0 in each of the EWCE-identified enriched cell types, were analysed independently for each brain region (striatum (n=26), frontal cortex (n=26), cerebellum (n=25)) using the STRING database. Including only interactions considered known within the database (derived from experimental data or curated databases), the same three networks of interactions were identified for each brain region. These included ADCY5, GNAL, GNB1, GNAO1, CACNA1A (network 1), TUBB4A, ACTB, YY1 (network 2), PRKRA, EIF2AK2 (network 3) (Figure 7)

**Figure 7.**
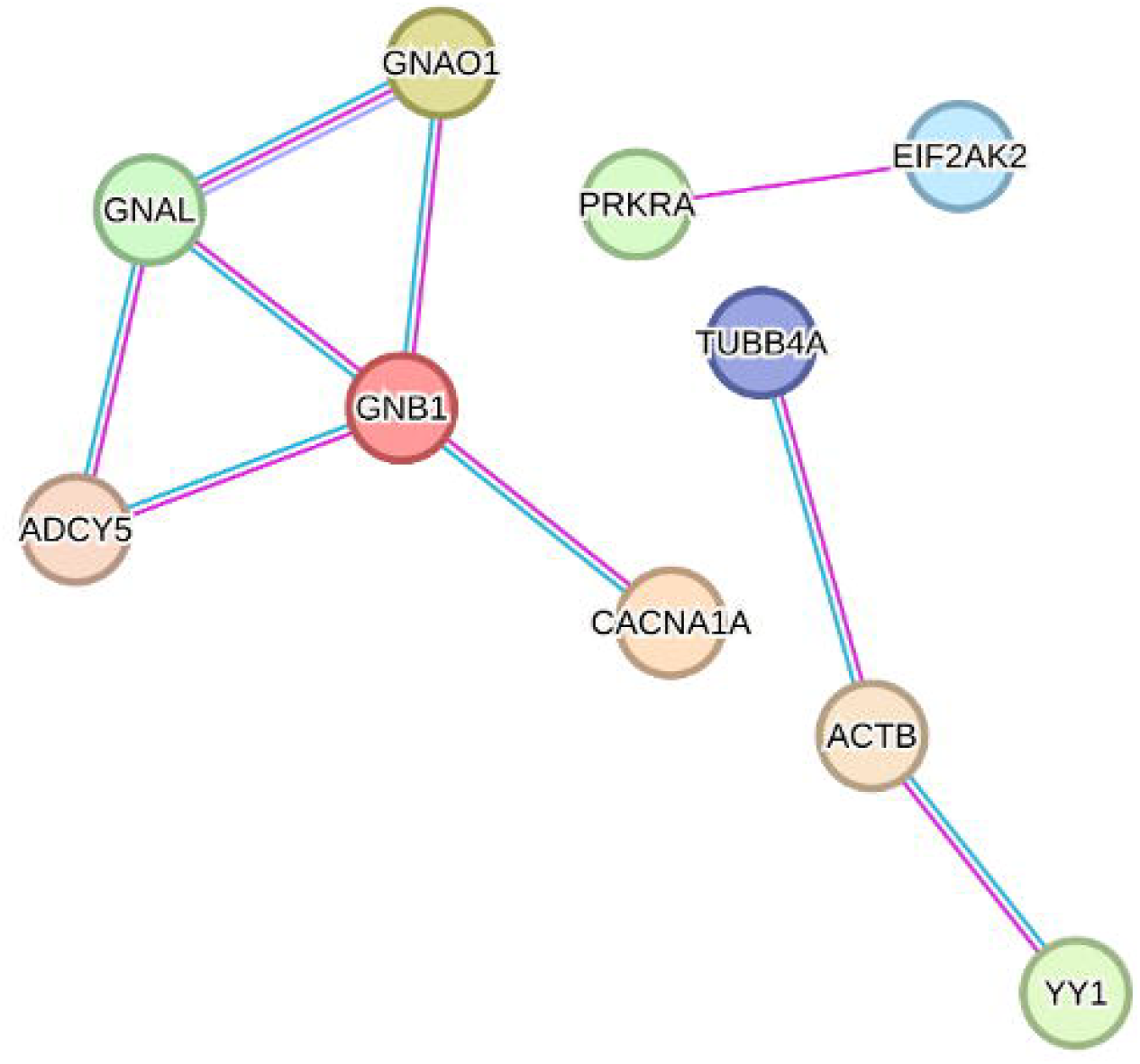
Dystonia gene expression specificity and protein-protein interaction analysis. Schematic representation of predicted protein-protein interactions involving dystonia genes expressed in each of the enriched cell populations across adult striatal, frontal cortex and cerebellar tissue. Nodes represent individual proteins, only interactions classified as known interactions included with blue links indicating data from curated databases and magenta links derived from experimentally determined data.

## Discussion

This study is the first to systematically evaluate the expression of a large group of genes causing monogenic forms of inherited dystonia in human brain tissue, spanning pre- and post-natal developmental periods. We found a notable increase in dystonia gene expression in the postnatal period, particularly in infancy and adulthood. Dystonia genes were most prominently expressed in the striatum, reinforced by significantly higher gene expression specificity. Cell type analysis showed predominant expression in neurons, notably inhibitory striatal MSNs, as well as excitatory and inhibitory neurons in the cerebral cortex and cerebellum. Gene Ontology analyses reinforced the role of dystonia genes in pathways linked to neuronal and synaptic structure and function, while predictive protein interaction analysis suggested interaction of the same 10 genes (*ADCY5, GNAL, GNB1, GNAO1, CACNA1A, PRKRA, EIF2AK2, TUBB4A, ACTB, YY1*) across all three brain regions.

Analysis of the BrainSpan dataset provided opportunity to investigate the developmental stages at which there were significantly higher levels of dystonia gene expression, potentially highlighting periods where deleterious variants may have the greatest impact. Higher expression was observed across all four post-natal stages, with largest effects observed in infancy and adulthood. Early motor features of dystonia, such as hypotonia, impaired co-ordination and gait difficulties, often emerge in infancy and have been linked to many of the genes included in this study.^3,28^ By contrast, adult-onset forms of dystonia are typically considered idiopathic. However, several recently discovered dystonia genes, such as *ANO3* and *GNAL*, show symptom onset in the 4^th^ and 5^th^ decade of life, suggesting the potential need for greater genetic testing in adult-onset forms of dystonia.^29^ Although statistically significant, the increased levels of dystonia gene expression in childhood and adolescence were lower than the other post-natal developmental periods, despite these being ages at which dystonia onset due to monogenic forms is frequently observed. While unexpected, this may be due to the delayed neurodevelopmental impact of deleterious variants in infancy with subsequent motor manifestation in childhood and adolescence.

The 44 dystonia genes were collectively enriched for higher expression in the striatum and cerebral cortex. These findings are in keeping with evidence across clinical studies and multiple model systems which have highlighted disruption to cortico-striatal networks, including increases to long-term potentiation (LTP) and reduced long-term depression (LTD) at cortico-striatal synapses as being central in dystonia pathogenesis.^30,31^ Although not frequently highlighted as regions of disruption in dystonia, additional enrichment of dystonia gene expression in amygdala and hippocampus may be relevant to non-motor symptoms of the dystonia phenotype, and in particular disruption to mood and memory.^6,32,33^ Finally, although not highlighted as a region with generally increased dystonia gene expression in this study (Figure 1B & 1C), several genes demonstrated high expression in the cerebellum (*CACNA1A, KCNMA1, AOPEP, GNAO1, GNAL, SGCE*), a brain region increasingly considered to be of importance in dystonia pathogenesis, and hypothesised to have a modulatory component within cortico-striato-thalamo-cerebellar circuits.^34,35^

Evaluation of single-nuclei transcriptomic data from the adult human striatum, frontal cortex and cerebellum revealed generally stronger expression of dystonia genes in neurons. This is consistent with findings from previous studies where disruption to excitatory-inhibitory balance is considered a key mechanism in dystonia pathophysiology with *in vivo* electrophysiological studies suggesting a reduction in inhibitory signalling may play a key role,^36-38^ while more recent work has suggested a more complex picture where both excess excitation and reduced inhibition contribute to the overall phenotype.^16,39^ Most genes with higher expression levels in neuronal cells were consistently expressed in both excitatory and inhibitory populations. Exceptions included *GNAL*, which was primarily expressed in inhibitory neurons of the frontal cortex, and *ANO3, KCTD17, MECR and RHOBTB2*, which showed predominant expression in excitatory neurons in this region. *GNAL* encodes the α-subunit of the G-protein, G_olf_, which is coupled to D1R and A2A adenosine receptors and activates type 5 adenylyl cyclase (AC5), encoded by another dystonia gene, *ADCY5*,^*40*^. This cell-type specificity, along with the functional links between these genes, will be valuable in designing disease-related models, such as those in patient derived stem cell and animal models.

Gene Ontology analysis demonstrated enrichment of dystonia genes in pathways involving neuronal structure (*Axon, Neuron projection, Main Axon, Somatodendritic compartment*), synapse (*Presynapse*), and neuronal signalling and function (*Axon Initial Segment* and *Membrane Protein Complex, Heterotrimeric G Protein Complex*). Axonal and neurite structural abnormalities have been identified across several neurodevelopmental disorders and are influenced by factors including neuronal activity, cell surface receptors and regulators of the actin cytoskeleton.^41-43^ Gene expression studies in *Thap1* (DYT6) murine models have identified neurite development as an enriched pathway across multiple brain regions, with signalling by the Rho family of GTPases, regulators of the actin cytoskeleton, emerging as a key linked pathway.^44,45^ Patient-derived induced pluripotent stem cell models harbouring *SGCE* mutations also demonstrated changes to dendritic morphology, including a higher number of branches, longer branches, and a more complex branching pattern, compared to their isogenic controls.^16^ The axon initial segment is a specialised membrane region from which action potentials are generated, and receives exclusive synaptic input from inhibitory GABAergic interneurons, neuronal subtypes already implicated in dystonia pathophysiology.^39,66^ Key to this role is the high density clustering of voltage-gated ion channels within the region, including Na^+^ (Nav1.1, 1.2, 1.6) and K^+^ (Kv1, 2.2) channels.^46,47^ More recently, patient derived stem cell models of two dystonia causing genes (*PRRT2* and *SGCE)* identified longer axon initial segments with each associated with a hyperexcitable neuronal phenotype, with *PRRT2* also found to be a negative regulator of Nav1.2 and Nav1.6 channels.^16,48^

Albeit at a lower level, dystonia genes were also found to be expressed in non-neuronal cells in the adult striatum, frontal cortex and cerebellum. While few studies have examined this to date, there is early evidence for some components of dystonia pathogenesis are mediated through impaired function of non-neuronal cells. One such example is *THAP1*, where conditional deletion has been shown to slow oligodendrocyte maturation and thereby delaying neuronal myelination in cell autonomous fashion. This is thought to be due to *THAP1* mutations disrupting a core set of genes involved in oligodendrocyte maturation, reducing DNA occupancy of *YY1*, a transcription factor required for oligodendrocyte maturation and another dystonia gene included in this analysis.^49^ *ATP1A3*, mutations in which cause rapid-onset dystonia parkinsonism, is known to have multiple brain isoforms, one of which is almost exclusively expressed in astrocytes. Further immunohistochemical post-mortem analyses have also identified the presence of astrocytes distributed across the striatum, cerebral cortex and cerebellum, again suggesting a potential role in pathogenesis.^50^

This study is the first to explore the expression of dystonia-causing genes across the human lifespan and multiple brain regions, highlighting central roles for the striatum, cerebral cortex and cerebellum and establishing the postnatal period as a critical stage in dystonia gene expression. The findings reveal consistent neuronal expression, involving both excitatory and inhibitory neurons, and implicate axons and synapses as focal points in dystonia pathophysiology. These developments pave the way for more targeted disease models, focused on specific brain regions and developmental windows. As the final clinical outcome is to identify novel small molecule and / or gene therapies for monogenic forms of dystonia, knowledge of dystonia gene expression patterns, such as that reported here, can help optimise the timing of developmental stage-specific therapies, and the ideal brain regions to target should these therapies require localised delivery.

## Acknowledgements

NEC is supported by grant MR/W017156/1 from the UK Medical Research Council. DC and NJB were supported by MRC project grants MR/T002379/1 and MR/Y003756/1. KJP is funded by an MRC Clinician-Scientist Fellowship & Transition Award (MR/P008593/1, MR/V036084/1).

## Author Contributions

(1) Conception and design of the study (DC, NEC, NJB, KJP), (2) acquisition and analysis of data (DC, DCF, PH, NJB, KJP), (3) drafting a significant portion of the manuscript or figures (DC, NJB, KJP), (4) manuscript editing and revision (DC, NEC, DCF, PH, NJB, KJP).

## Data Availability

Scripts for all analyses described are available at https://github.com/Dazcam/dystonia_snRNAseq_2024

