## Supplementary Material for "Developmental, neuroanatomical and cellular expression of genes causing dystonia"

**Supplementary Table 1: Dystonia phenotype associated genes**

| Gene (OMIM, DYT) | Chromosome (start location; end location) | Inheritance | Clinical Characteristics |
|---|---|---|---|
| ***SLC2A1*** (601042, DYT9] | chr1 (42925353; 42958868) | AD | Childhood onset. Paroxysmal choreoathetosis and progressive spastic paraplegia. Cognitive Impairment. |
| ***HPCA*** (224500, DYT2) | chr1 (32886491; 32894646) | AR | Childhood- or adolescent-onset. Distal limb dystonia, progresses to involve neck, orofacial and cranio-cervical regions |
| ***COX20*** (619054) | chr1 (244835306; 244845063) | AR | Associated with early-onset hypotonia, dystonia, ataxia, sensory neuropathy and dysarthria |
| ***MECR*** (617282, DYT29) | Chr1 (29167696; 29230934) | AR | Childhood-onset dystonia, optic atrophy and basal ganglia abnormalities with varying phenotypic severity |
| ***GNB1*** (616973) | Chr1 (1785286; 1891087) | AD | Global developmental delay, cognitive impairment, hypotonia, dystonia, seizures, poor growth |
| ***SPR*** (612716, DYT5) | chr2 (72887408; 72892158) | AR | L-dopa responsive, diurnally fluctuating movement disorder, usually associated with cognitive delay and severe neurological dysfunction |
| ***EIF2AK2*** (619687, DYT3) | chr2 (37099210; 37157192) | AD, AR | Focal or generalised dystonia in first 2 decades of life. Slow progression, potentially leading to difficulties with mobility, dysarthria, or dysphagia. Incomplete penetrance and intrafamilial phenotypic heterogeneity. |
| ***PRKRA*** (612067, DYT16) | chr2 (178431414; 178451175) | AR | Childhood- or adolescent-onset of limb dystonia with parkinsonism, commonly with cranio-cervical or laryngeal dystonia with progression to generalised involvement |
| ***PNKD*** (118800) | chr2 (218270514; 218346784) | AD | Attacks of dystonia, chorea and athetosis. Attacks may be precipitated by stress, fatigue, caffeine, alcohol, menstruation, and may last minutes to hours. |
| ***ADCY5*** (606703) | chr3 (123282296; 123449090) | AD, AR | involves choreiform, myoclonic and dystonic movements involving neck, limb, and facial muscles in 1st decade of life |
| ***SLC6A3*** (613135, PKDYS1) | Chr5 (1392794; 1445440) | AR | Infantile-onset parkinsonism-dystonia-1 (PKDYS1) associated with orolingual and limb dyskinesia, dystonia, chorea or parkinsonian features. |
| ***SQSTM1*** (617145) | Chr5 (179806393; 179838078) | AR | Childhood-onset neurodegeneration with ataxia, dystonia, gaze palsy and cognitive impairment. |
| ***SERAC*** (614739) | Chr6 (158109519; 158168262) | AR | Childhood-onset delayed psychomotor development, dystonia, sensorineural deafness. Cerebral and cerebellar atrophy, basal ganglia lesions on brain imaging. |
| ***SGCE*** (159900, DYT11) | chr7 (94584980; 94656133) | AD | Upper body myoclonus and dystonia |
| ***DDC*** (608643) | chr7 (50458442; 50565405) | AR | Onset in infancy and early childhood with hypotonia, oculogyric crises and dystonia |
| ***ACTB*** (607371) | Chr7 (5527148; 5530601) | AD | Congenital or childhood-onset sensorineural deafness, subsequent dystonia with frequent bulbar involvement. Skeletal abnormalities, dysmorphic features and developmental delay observed in some |
| ***THAP1*** (602629, DYT6) | chr8 (42836674; 42843325) | AD | Onset typically in the second decade of life with early involvement of craniofacial muscles with secondary generalisation often involving the arms and larynx. |
| ***RHOBTB2*** (618004) | Chr 8 (22950813; 23020199) | AD | Infancy-onset seizures, cognitive and motor impairment including dystonia, dysarthria or aphasia, hypotonia and dysmorphic features. |

| | | | |
|---|---|---|---|
| ***TOR1A***<br>(128100, DYT1) | chr9<br>(129812942; 129824136) | AD | Onset in first two decades of life frequently beginning in the lower limbs and progressing from isolated to generalised involvement. |
| ***AOPEP***<br>(619565, DYT31) | Chr9<br>(94726699; 95150224) | AR | Dystonia involving the upper and lower limbs, craniofacial region and trunk with onset from childhood to young adult life. Associated orofacial dyskinesia resulting in dysphagia reported in some cases. |
| ***KCNMA1***<br>(609446) | chr10<br>(76884869; 77637902) | AD | Paroxysmal Non-Kinesigenic Dyskinesia |
| ***DNAJC12***<br>(617384) | chr10<br>(67796669; 67838188) | AR | Mild non-BH4-deficient Hyperphenylalaninaemia - characterised by increased serum phenylalanine. Juvenile or young-adult-onset nonprogressive dopa-responsive parkinsonism |
| ***ANO3***<br>(615034, DYT24) | chr11<br>(26332130; 26663289) | AD | Infantile- or adult-onset segmental dystonia, commonly involving the cervical, laryngeal and upper limb muscles. |
| ***TH***<br>(605407, DYT5) | chr11<br>(2163929; 2171815) | AR | Infancy onset dopa-responsive dystonia |
| ***TMEM151A***<br>(620245) | Chr11<br>(66291894; 66296664) | AD | Childhood-onset paroxysmal kinesigenic dyskinesia characterised by dystonia, chorea, athetosis and other hyperkinetic movement disorders typically triggered by sudden movement or stress |
| ***SCN8A***<br>(617080) | chr12<br>(51591233; 51812864) | AD | Paroxysmal kinesigenic dyskinesia, benign infantile seizures. |
| ***KCNA1***<br>(160120) | Chr12<br>(4909905; 4918256) | AD | Paroxysmal kinesigenic dyskinesia, precipitated by sudden movements or prolonged activity. |
| ***GCH1***<br>(128230, DYT5) | chr14<br>(54842008; 54902824) | AD, AR | Childhood-onset dystonia, typically of onset in the lower limbs, diurnal fluctuation and improves with levodopa treatment. May progress to segmental or generalised involvement. |
| ***FOXG1***<br>(613454) | Chr14<br>(28766787; 28770277) | AD | Severe developmental delay, microcephaly, epilepsy, dystonia, chorea and orolingual dyskinesias |
| ***YY1***<br>***(617557)*** | Chr14<br>(100239144; 100282788) | AD | Infancy-onset psychomotor developmental delay, cognitive impairment, dystonia, dysmorphic facial features and feeding difficulties |
| ***PRRT2***<br>(128200) | chr16<br>(29812193; 29815881) | AD | Episodic kinesigenic dyskinesia - characterised by recurrent and brief attacks of involuntary movements triggered by sudden voluntary movement. Onset childhood or early adulthood and can involve dystonic posture, chorea or athetosis. Symptoms respond to carbamazepine or phenytoin. |
| ***GNAO1***<br>(617493) | chr16<br>(55991504; 56162084) | AD | Infantile or Childhood-onset segmental or generalised dystonia prominently affecting the upper body. Chorea and athetosis may also occur. |
| ***VAC14***<br>(617054) | Chr16<br>(70687439; 70801158)) | AR | Motor developmental milestone regression during 1st year of life including dystonia, speech and gait impairment. Evidence of nigrostriatal degeneration on brain imaging. |
| ***TSPOAP1***<br>(620456, DYT22) | Chr17<br>(58301231; 58328795) | AR | Focal dystonia, tremor and mild cognitive impairment |
| ***GNAL***<br>(615073, DYT25] | chr18<br>(11752085; 11885685) | | Adult-onset focal dystonia usually involving the neck. Dystonia most often progresses to involve other regions, particularly face and laryngeal muscles, and less commonly trunk and limbs. |

| | | | |
|---|---|---|---|
| ***TUBB4A*** (128101, DYT4) | chr19 (6494319; 6502309) | AD | Whispering dysphonia. Onset in 2nd-3rd decade of life with progressive laryngeal dysphonia followed by involvement of other body parts such as neck or limbs. |
| ***KMT2B*** (617284, DYT28) | chr19 (35727156; 35728171) | AD | Progressive dystonia in first decade of life. Usually begins focally in the lower limbs, progression to other body regions, including upper limbs, neck and orofacial regions |
| ***ATP1A3*** (128235, DYT12) | chr19 (41966584; 41994270) | AD | Rapid-onset dystonia-parkinsonism - abrupt onset asymmetric dystonia and parkinsonism in young adulthood, often after a trigger such as physical overexertion, trauma, heat or fever. |
| ***CACNA1A*** (617106) | Chr19 (13206442; 13506479) | AD | Early life onset of seizures, global developmental delay, severe intellectual impairment, axial hypotonia, peripheral hypertonia, hyperreflexia, tremor, dystonia, ataxia. |
| ***VPS16*** (619291, DYT30) | chr20 (2840745; 2866732) | AD | Onset of symptoms in first decades of life with oromandibular, cervical, bulbar or UL dystonia, with slow progression to generalised dystonia |
| ***FITM2*** (618635) | Chr20 (44302840; 44311202) | AR | Global developmental delay, dystonia, progressive sensorineural hearing loss, motor skill regression, reduced growth and low body mass index. |
| ***KCTD17*** (616398, DYT26) | chr22 (37051742; 37063390) | AD | Myoclonic jerks affecting UL in 1st/2nd decade of life. Disorder is progressive with patients later developing dystonia with prominent involvement of the cranio-cervical regions and sometimes trunk and/or lower limbs. |
| ***TAF1*** (314250, DYT3) | Chr X (71366357; 71530525) | XLR | Onset with initial focal dystonia in the lower limbs or oromandibular regions, progressing to generalized involvement with subsequent development of parkinsonian features. |
| ***BCAP31*** (300475) | Chr X (153700492; 153724387) | XLR | Early-onset failure of psychomotor development, dystonia, dysmorphic facial features, sensorineural deafness, cerebral hypomyelination. |
| ***TIMM8A*** (304700) | Chr X (101345661; 101348742) | XLR | Childhood-onset progressive sensorineural deafness, dystonia, cognitive impairment, cortical blindness and psychiatric symptoms. |

**Legend:** AD: Autosomal Dominant, AR: Autosomal Recessive, XLR: X-Linked Recessive

**Supplementary Table 2: Definition of Developmental Stages and Sample Numbers included from each dataset.**

| | BrainSpan (n) | | | | | | | |
|---|---|---|---|---|---|---|---|---|
| **Developmental Stage** | **Prefrontal Cortex** (Total = 112) | **Non-Prefrontal Cortex** (Total = 145) | **Primary Motor-Sensory Cortex** (Total = 48) | **Striatum** (Total = 31) | **Thalamus** (Total = 26) | **Cerebellum** (Total = 30) | **Hippocampus** (Total = 28) | **Amygdala** (Total = 27) |
| **Early Fetal** (<13 pcw) | 30 | 32 | 15 | 12 | 6 | 6 | 8 | 8 |
| **Midfetal** (13-24 pcw) | 18 | 27 | 7 | 6 | 5 | 4 | 5 | 4 |
| **Late Fetal** (24 pcw-0 months) | 5 | 8 | 2 | 1 | 1 | 1 | 1 | 1 |
| **Infancy** (0 months -1 year) | 8 | 12 | 3 | 2 | 3 | 3 | 2 | 2 |
| **Childhood** (1-13 years) | 23 | 29 | 7 | 4 | 4 | 8 | 5 | 4 |
| **Adolescence** (13-20 years) | 7 | 8 | 2 | 1 | 1 | 2 | 1 | 2 |
| **Adulthood** (>20 years) | 21 | 29 | 12 | 5 | 6 | 6 | 6 | 6 |

**<u>Supplementary Table 3: Summary of scRNA-seq data downloaded from Siletti et al</u>**

| Region | Dissection | Abbreviation | Download link |
|---|---|---|---|
| Striatum | Basal nuclei (BN) - Body of the Caudate - CaB | CaB | a48ed83c-7db8-4419-82f1-7100aefb3cfe |
| | Basal nuclei (BN) - Putamen - Pu | Pu | b92375fd-dafe-44c6-8523-f78c73660e85 |
| Cerebellum | Cerebellum (CB) - Cerebellar deep nuclei - CbDN | CbDN | dd24ac0c-07c1-4192-b774-7b7ac146f5e6 |
| | Cerebellum (CB) - Cerebellar Vermis - CBV | CBV | db5392a6-8e1e-4ca7-8dd9-137c536f11d8 |
| | Cerebellum (CB) - Lateral hemisphere of cerebellum - CBL | CBL | 4ff30bec-1b28-4d36-b7c2-fcb9f07919b4 |
| Frontal Cortex | Cerebral cortex (Cx) - Frontal agranular insular cortex - FI | FI | 215cec1c-1ec4-4c1d-ba73-ce709ac6382f |
| | Cerebral cortex (Cx) - Gyrus rectus (ReG) - Medial orbitofrontal cortex - A14 | A14 | 0cfc9ba4-b520-4bf3-8a1d-5cc0d9eae617 |
| | Cerebral cortex (Cx) - Inferior frontal gyrus (IFG) - Ventrolateral prefrontal cortex - A44-A45 | A44-A45 | 4fd62107-01a2-4e3f-a374-874a2078c30c |
| | Cerebral cortex (Cx) - Middle frontal gyrus (MFG) - A46 | A46 | 8f513d63-f891-4022-a9e5-687bb1767050 |
| | Cerebral cortex (Cx) - Posterior intermediate orbital gyrus (POrG) - Caudal division of OFCi - A13 | A13 | 00724105-4ab6-4e0f-abf9-33825f3b2206 |
| | Cerebral cortex (Cx) - Precentral gyrus (PrCG) - Primary motor cortex - M1C | M1C | 209fa352-40c5-49d4-b829-29847390ad1c |
| | Cerebral cortex (Cx) - Rostral gyrus (RoG) - Dorsal division of MFC - A32 | A32 | e578d6fe-215a-4123-9350-030da4a83457 |
| | Cerebral cortex (Cx) - Subcallosal Gyrus (SCG) - Subgenual (subcallosal) division of MFC - A25 | A25 | abb33669-3610-4cce-ae94-07f06b99a099 |

**Supplementary Table 4: Summary of BrainSpan developmental stage linear model analysis**

| Developmental Stage | Estimate | Standard Error | P-value | FDR |
|---|---|---|---|---|
| Early Fetal | -0.36361191 | 0.01443276 | 9.14E-138 | **6.40E-137** |
| Mid Fetal | -0.00367720 | 0.01708975 | 0.83 | 0.83 |
| Late Fetal | 0.03112917 | 0.03103658 | 0.32 | 0.37 |
| Infancy | 0.20345055 | 0.02384928 | 1.56E-17 | **3.65E-17** |
| Childhood | 0.04729065 | 0.01647073 | 4.09E-01 | **5.73E-03** |
| Adolescence | 0.14368405 | 0.02846382 | 4.51E-07 | **7.89E-07** |
| Adulthood | 0.24442763 | 0.01584744 | 2.41E-53 | **8.43E-53** |

**Legend:** Bold highlights statistically significant results

**Supplementary Table 5: Adult striatal tissue single cell RNA-seq data. Median expression values for individual genes across neuronal and non-neuronal cell types**

**Supplementary Table 6: Adult frontal cortical tissue single cell RNA-seq data. Median expression values for individual genes across neuronal and non-neuronal cell types**

**Supplementary Table 7: Adult cerebellar tissue single cell RNA-seq data. Median expression values for individual genes across neuronal and non-neuronal cell types**

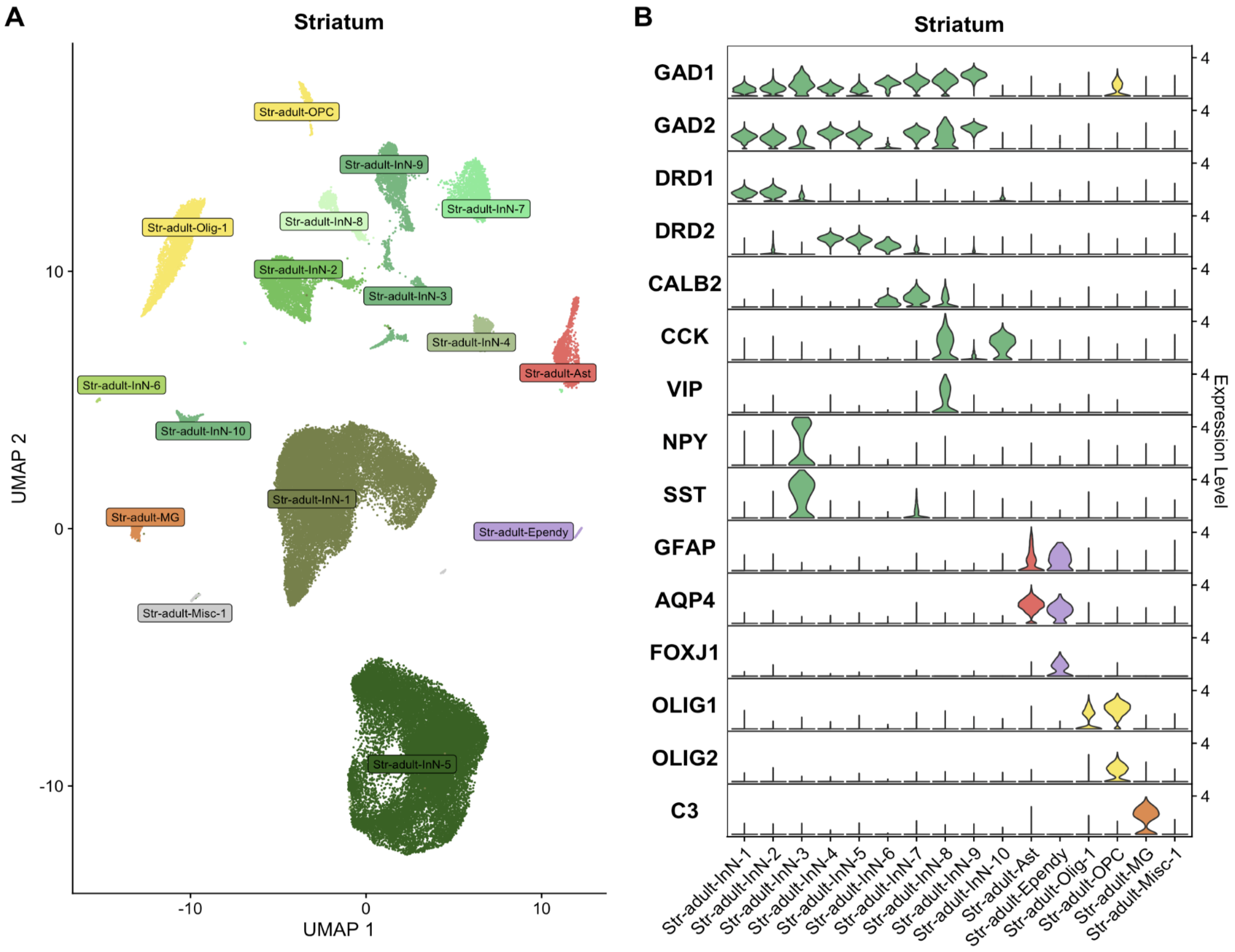


**Supplementary Figure 1. Cell populations of the adult human striatum analysed in this study.** Single nuclei RNA sequencing data generated from the striatum in the study of Siletti et al (*Siletti et al, Science 2023 Oct 13;382(6667):eadd7046*) were downloaded and analysed using Seurat 5.0.2. A) UMAP showing clusters of striatal nuclei based on gene expression. B) Violin plots showing expression of cell type-specific markers as the basis of cell annotations. Ast: Astrocyte, Ependy: Ependymal cells, InN: inhibitory neuron, MG: microglia, Misc: Miscellaneous, Olig: Oligodendrocyte, OPC: oligodendrocyte precursor cell.

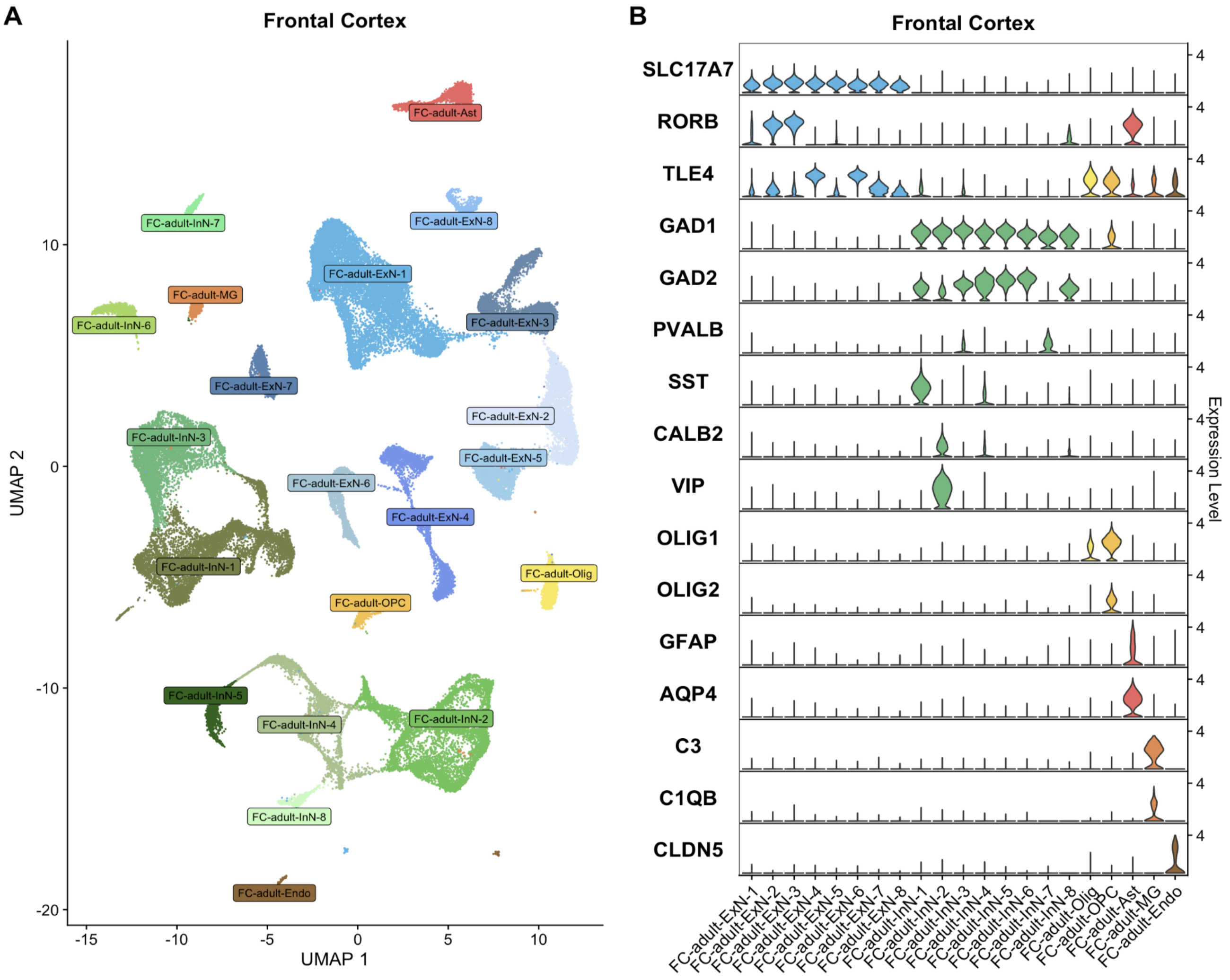


**Supplementary Figure 2. Cell populations of the adult human frontal cortex analysed in this study.** Single nuclei RNA sequencing data generated from the frontal cortex in the study of Siletti et al (*Siletti et al, Science 2023 Oct 13;382(6667):eadd7046*) were downloaded and analysed using Seurat 5.0.2. A) UMAP showing clusters of striatal nuclei based on gene expression. B) Violin plots showing expression of cell type-specific markers as the basis of cell annotations. Ast: Astrocytes, Endo: Endothelial cells, ExN: excitatory neuron, InN: inhibitory neuron, MG: microglia, Olig: Oligodendrocyte, OPC: oligodendrocyte precursor cell.

**Supplementary Figure 3**

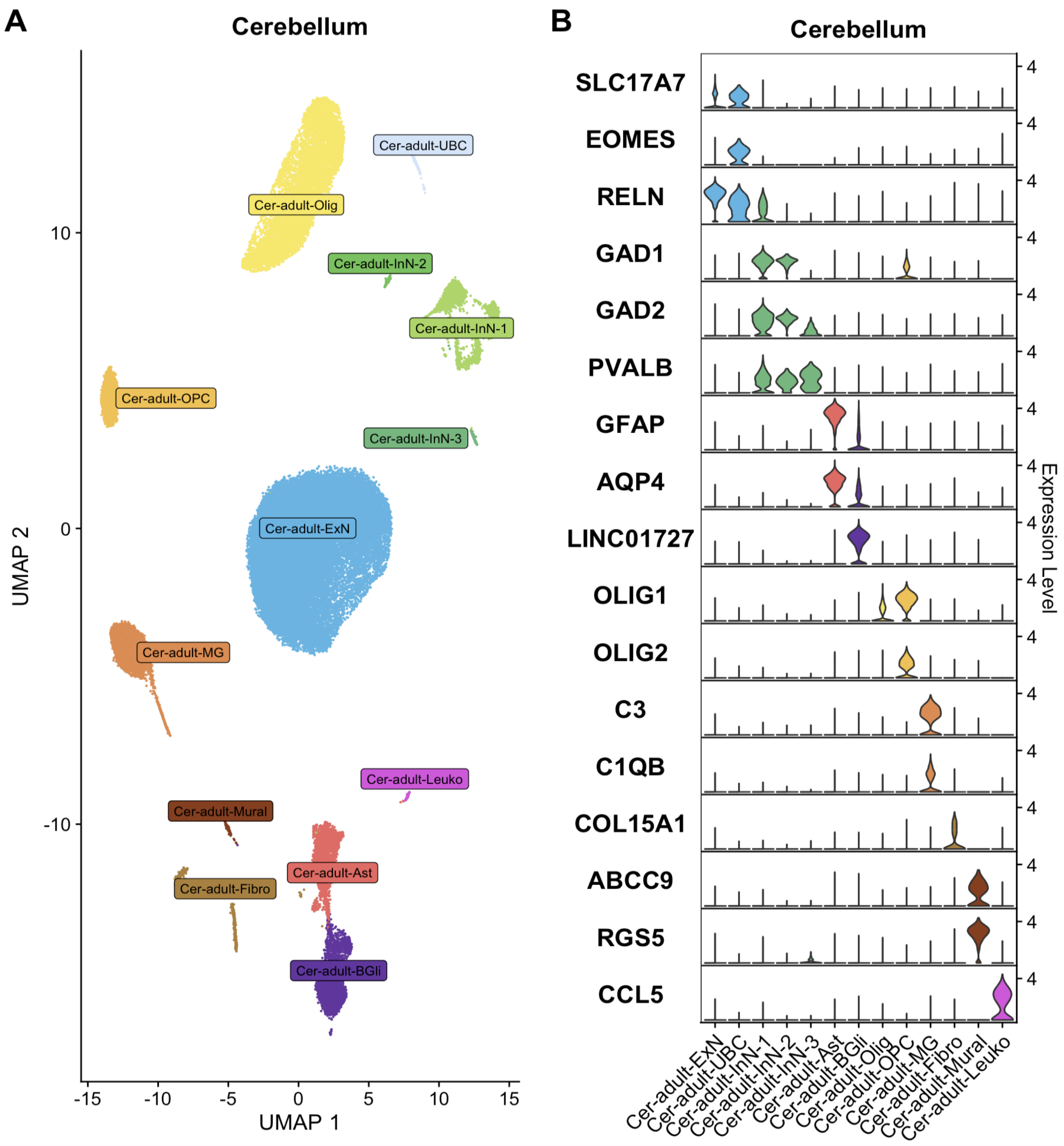


**Supplementary Figure 3. Cell populations of the adult human cerebellum analysed in this study.** Single nuclei RNA sequencing data generated from the cerebellum in the study of Siletti et al (*Siletti et al, Science 2023 Oct 13;382(6667):eadd7046*) were downloaded and analysed using Seurat 5.0.2. A) UMAP showing clusters of striatal nuclei based on gene expression. B) Violin plots showing expression of cell type-specific markers as the basis of cell annotations. Ast: Astrocytes, BGli: Bergmann glial cells, Cer: Cerebellum, ExN: excitatory neuron, Fibro: Fibroblast, InN: inhibitory neuron, Leuko: Leukocyte, Mural: Mural cells, MG: microglia, Olig: Oligodendrocyte, OPC: oligodendrocyte precursor cell, UBC: Unipolar Brush cells.

**Supplementary Figure 4: Cellular expression of individual dystonia genes in fetal ganglionic eminence, frontal cortex and cerebellar tissue**

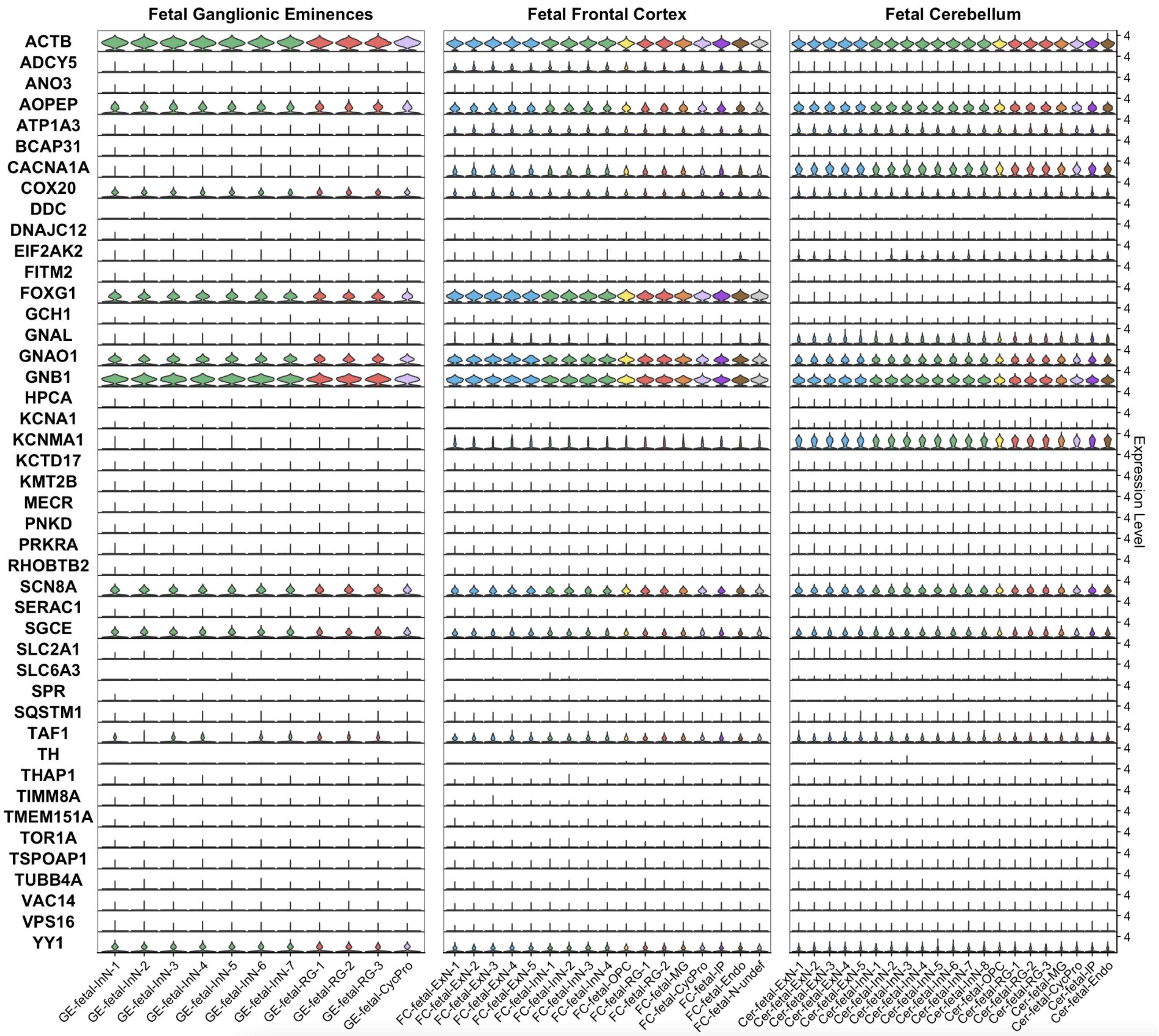


**Supplementary Figure 4. Dystonia gene expression in cell populations of the human fetal ganglionic eminences, frontal cortex and cerebellum.** Data generated by Cameron et al (*Cameron et al, Biological Psychiatry 2023 Jan 15;93(2):157-166*) from 3 fetuses aged 14-15 post-conception weeks using single-nuclei RNA sequencing. Violin plots show expression of each gene in each cell population. CycPro: cycling progenitor cell, Endo: endothelial cell, ExN: excitatory neuron (developing form), InN: inhibitory neuron (developing form), IP: intermediate progenitor, MG: microglia, N-undef: neuron of undefined class, OPC: oligodendrocyte precursor cell, RG: radial glia.

**<u>Supplementary Figure 5:</u> EWCE analysis of fetal ganglionic eminence, frontal cortex and cerebellar tissue**

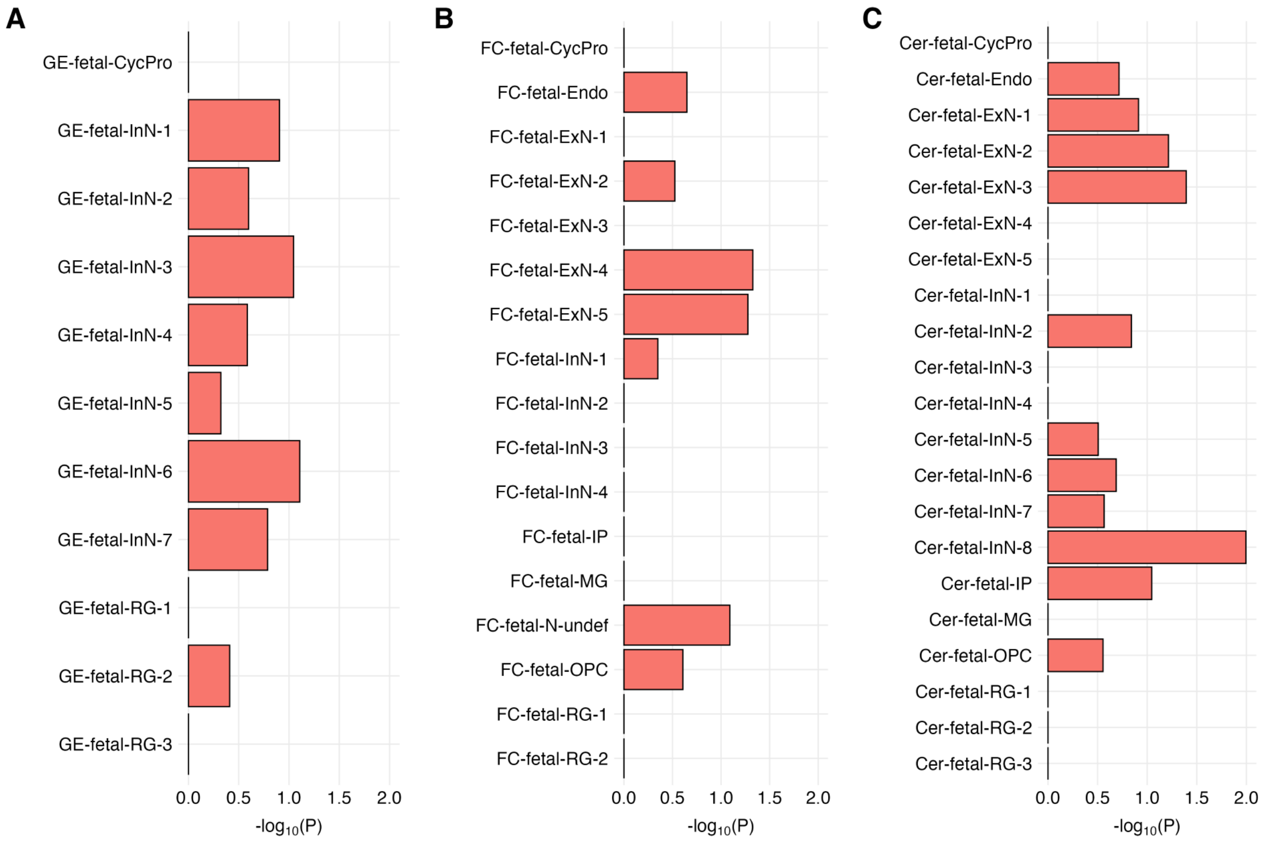


**<u>Legend:</u>** Bar plots demonstrating -log10P enrichment values for the 44 dystonia genes across individual cell subtypes in fetal ganglionic eminence, frontal cortex and cerebellar tissue (red bars). Statistical significance was determined using a false discovery rate (FDR) correction, with enrichments considered significant at PFDR < 0.05. No enrichment was significant at this PFDR. CycPro: cycling progenitor cell, Endo: endothelial cell, ExN: excitatory neuron (developing form), InN: inhibitory neuron (developing form), IP: intermediate progenitor, MG: microglia, N-undef: neuron of undefined class, OPC: oligodendrocyte precursor cell, RG: radial glia.

**Supplementary Figure 6: EWCE analysis of adult striatal, frontal cortex and cerebellar tissue**

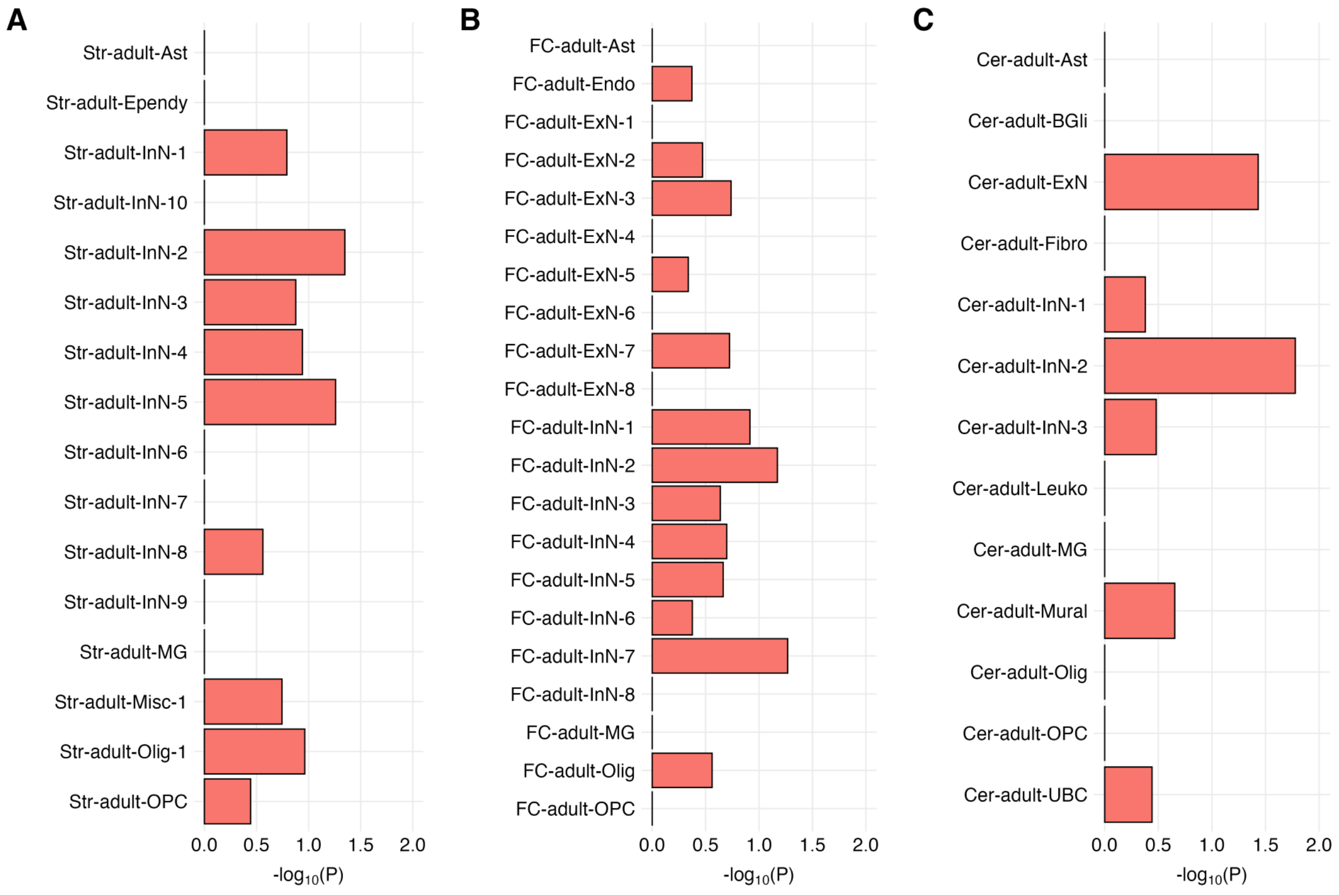


**Legend:** Bar plots demonstrating -log10P enrichment values for the 44 dystonia genes across individual cell subtypes in striatal (A), frontal cortex (B) and cerebellar tissue (C) (red bars). Statistical significance was determined using a false discovery rate (FDR) correction, with enrichments considered significant at PFDR < 0.05. No enrichment was significant at this PFDR. Ast: Astrocytes, BGli: Bergmann glial cells, Ependy: Ependymal cells, ExN: excitatory neuron, Fibro: Fibroblast, InN: inhibitory neuron, Leuko: Leukocyte, MG: microglia, Misc: Miscellaneous, Mural: Mural cells, Olig: Oligodendrocyte, OPC: oligodendrocyte precursor cell, UBC: Unipolar Brush cells.
